# Patterned alginate hydrogel spatially guides collagen fibrillogenesis, viscoelasticity and endothelial cell invasion

**DOI:** 10.64898/2026.09.02.748900

**Authors:** C.A. Garrido, D.S. Garske, B. Häßel, C. Bastard, J. Kamp, L. De Laporte, G.N. Duda, K. Schmidt-Bleek, A. Cipitria

## Abstract

Angiogenesis following injury has been shown to be driven by fibrillar proteins of the extracellular matrix (ECM), such as collagen. However, the use of protein-based biomaterials presents some challenges, such as uncontrolled degradation and limited tuneability. We demonstrate how to create patterned interpenetrating networks (IPNs) based on covalently crosslinked alginate and physically crosslinked collagen that provide suitable mechanical properties to support migration of endothelial cells (ECs) in a spatially controlled manner. Low molecular weight alginate is functionalized with norbornene (N) or tetrazine (T), which enables two independent covalent crosslinking methods: UV-mediated and degradable crosslinks with matrix metalloproteinase (MMP) sensitive peptides (Deg) and slower spontaneous N:T non-degradable crosslinks (noDeg). Using photolithography, patterns in degradation, collagen fibrillogenesis, microarchitecture and matrix viscoelasticity are created. The potential of such 3D patterned alginate-collagen (Alg-Col) IPNs to spatially guide EC invasion and proliferation was tested in a microfluidics platform resembling an early healing setting. Only regions combining collagen fibrillogenesis, alginate degradability and viscoelasticity demonstrated EC cell invasion similar to the ones found *in vivo* following injury. The 3D patterned Alg-Col IPNs are compatible with microfluidics, offer an strategy to widen the applications of protein-based hydrogels and present a versatile platform for tissue engineering and disease modeling.

**Table of Contents:** 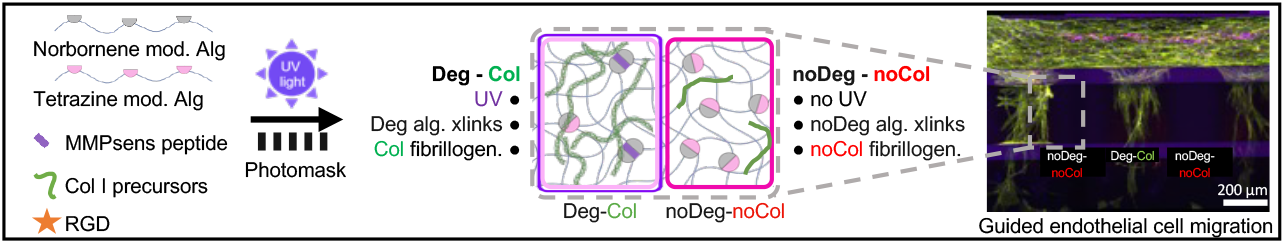

3D patterned alginate–collagen interpenetrating networks spatially control collagen fibrillogenesis, matrix degradability, microarchitecture, and viscoelasticity. By combining orthogonal crosslinking with photolithographic patterning, distinct matrix regions guide endothelial cell invasion and proliferation. These hybrid materials recapitulate key angiogenic responses following injury and can be readily incorporated into microfluidic chips, providing a versatile platform for tissue engineering and disease modeling.

## 1. Introduction

The development of new biomaterial systems has made important advances in understanding the role of the extracellular matrix (ECM) in actively directing cell behavior with applications in tissue engineering and disease modeling. Emerging approaches include dynamic biomaterials [1], stimuli-responsive materials [2], and microscale guiding structures [3], all of which highlight the potential of material design to guide cellular function. Collagen is the most abundant ECM protein [4] and biomaterials derived from it are promising candidates due to their biocompatibility [5], viscoelastic properties [6]and potential for structural modification [7]. The effect of spatial distribution of collagen to resemble its occurrence in nature has been broadly studied in 2D, showing the potential of collagen anisotropic distribution to control cell morphology and proliferation [8], as well as cell alignment [9]. However, mimicking tissues with 3D collagen anisotropy brings additional challenges as native collagen has limited mechanical properties, uncontrolled degradation and matrix contraction over time [10–12]. To overcome these limitations, collagen can be mixed with other polymers to create composites [13] and interpenetrating polymer networks (IPNs) [7].

IPNs are a broad class of polymer mixtures, classically defined as materials containing two or more networks of polymers with at least one being polymerized or crosslinked in the presence of the other [14]. Collagen can be mixed with other polymers, such as polyethylene glycol (PEG) to modify collagen’s microarchitecture and improve the mechanical properties [15,16], hyaluronic acid to further tune the porosity and closer mimic macromolecule transport in the ECM [17], polyurethane to increase the resistance to thermal and hydrolytic degradation [18], or even collagen itself by combining fibrillar-physically assembled collagen with covalently crosslinked collagen to combine cell-ECM interactions and structural integrity [16].

IPNs combining collagen and alginate have been broadly studied, as the inert nature and mechanical versatility of alginate help overcome the mechanical limitations of collagen [19], preserving the high biocompatibility of both materials [5] and allowing versatile applications, such as microsphere production [20] or bioprinting [21]. However, most of these studies describing alginate-collagen (Alg-Col) IPNs are based on ionically crosslinked alginate with calcium ions[22,23]. This type of IPNs have shown interesting properties, such as variable stiffness independent of the substrate architecture [23], tunable binding sites [24] and efficient drug delivery [25]. However, it has limited stability under physiological conditions due to the diffusion of the crosslinking calcium ions [26–29]. Therefore, a strategy is required to preserve the advantages of the alginate network but overcoming the limitations of ionic crosslinking.

Covalently crosslinked alginate offers an opportunity to control mechanical properties, degradation and biological functionalization through different chemical reactions (e.g. thiol- ene [30] or click chemistries [31]). Consequently, covalently crosslinked alginate has been used for cell encapsulation [27], controlled drug [32] or cell delivery [33], and disease modeling [34]. Alg-Col IPNs involving sequential crosslinking methods of alginate (covalent and ionic crosslinking) have shown to modulate the viscoelastic properties of the matrix and influence the immune response of human stromal cells [35]. Furthermore, covalently crosslinked alginate can be patterned through photolithography, with stiffness and adhesion patterns for 2D cell growth [36] or with stiffness and degradation patterns for 3D cell growth [15,37,38]. Despite the advantages of covalently crosslinked alginate, the elastic behavior of the materials and its derivates limits its soft tissue mimicking applications.

ECM viscoelastic properties can promote endothelial cell (EC) migration [39] while collagen fiber density and alignment can guide angiogenesis [40]. For this reason, ionically crosslinked Alg-Col IPNs have been used in vasculogenesis, as they reduce protein-based biomaterial contraction [41], support the co-culture of human mesenchymal stromal cells (hMSCs) and ECs [42], and enhance ECs invasion [43]. On the other hand, anisotropic materials have shown promising results guiding ECs, such as enhanced sprouting [44] and cell arrangement [45]. Yet, there are no reports on 3D patterned Alg-Col IPNs for angiogenesis research.

Furthermore, combining biomaterials with microfluidics is a promising approach to study angiogenesis as it provides control over shear stress, spatial architecture, nutrient, and chemical transport properties [46]. 3D culture of EC cells in microfluidic chips has allowed to replicate multiple cell niches, such as lymphatic vessels [47], the blood brain barrier [48] or the breast cancer microenvironment in disease models [49]. These dynamic 3D models allow for drug or growth factor screening [50], EC interaction with other cell types [51,52], and most importantly, incorporate ECM properties supporting angiogenesis [46].

In this study, we present 3D patterned Alg-Col IPNs with covalently crosslinked alginate and physically crosslinked collagen that allow for spatial patterning of 3D collagen fibrillogenesis, collagen fiber network formation, and matrix viscoelasticity, rendering the system anisotropic. This was achieved by photolithography using chemically modified alginate that forms degradable regions when exposed to UV light, and non-degradable regions in the unexposed areas. The 3D patterned IPNs were incorporated in microfluidic devices and showed that regions with fibrillar collagen architecture and viscoelastic properties support patterned EC invasion. This technology provides a versatile platform for tissue engineering and disease modeling and opens many opportunities to create organs-on-a-chip or larger tissue models using bioprinting.

## 2. Materials and methods

### 2.1. Alginate modification

The basis of the IPNs is collagen and modified alginate. Collagen type I precursor solution (FibriCol, #5133, Biomatrix) and low molecular weight sodium alginate with high guluronic acid (MW 75 kDa Pronova UP VLVG; #4200501, NovaMatrix) were used.

The alginate was functionalized with norbornene or tetrazine through carbodiimide chemistry. The coupling of norbornene (N, TCI Chemicals, #N0907) or tetrazine (T, Conju-probe, #CP-6021) to the alginate backbone was performed as previously described, adapting the modification rates to the lower molecular weight of VLVG [37,38]. Alginate modification was performed with a theoretical degree of substitution (DS_theo_) of 500 for norbornene and DS_theo_ 170 for tetrazine. The two chemical groups were not coupled to the same alginate molecules. To ensure appropriate norbornene to tetrazine (N:T) ratios for crosslinking, the actual DS (DS_actual_) was determined by nuclear magnetic resonance (NMR) measurements, using a 1.5% w/v alginate solution in deuterium oxide (64 scans; Agilent 400 MHz Premium COMPACT equipped with Agilent OneNMR Probe) and analyzed using MestreNova Software (version 14.6). Representative NMR spectra can be found in Suppl. Figure S1 and the DS are summarized in Suppl. Table S1.

### 2.2. Production of degradable or non-degradable bulk IPNs with covalent-alginate and physical-collagen crosslinking

Initially, bulk 3D IPNs with covalent-alginate and physical-collagen crosslinking (Alg-Col IPN) were fabricated without a photomask based on previously reports, where a covalently crosslinked alginate was produced [36–38]. The following sections describe in detail the fabrication of Alg-Col IPNs. Briefly, the protocol was adapted from previously published work [36–38] to enable collagen fibrillogenesis by replacing PBS with HEPES buffer (#15630106, Gibco), which provided improved pH control. In addition, the photoinitiator Irgacure 2959 was substituted with lithium phenyl-2,4,6-trimethylbenzoylphosphinate (LAP, #900889, Sigma), owing to its higher reaction efficiency (1.5 versus 10 min) and reduced phototoxicity to cells. A schematic overview of the materials described in the following sections is provided in Figure 1. The precursor solution was the same for both the non-degradable and degradable Alg-IPN to later allow for patterning. For the non-degradable Alg-IPN, the solution is not exposed to UV light and crosslinks via spontaneous norbornene-tetrazine click reaction. For the degradable Alg-IPN, the solution is exposed to UV light at 5.7 mW/cm^2^ for 1.5 min, favoring the thiol-ene click reaction between the norbornene-modified alginate (N-Alg) and the MMP-sensitive (MMPsens) peptide (GCRD-VPMS ↓ MRGG-DRCG, 95 % purity; WatsonBio).

**Figure 1:**
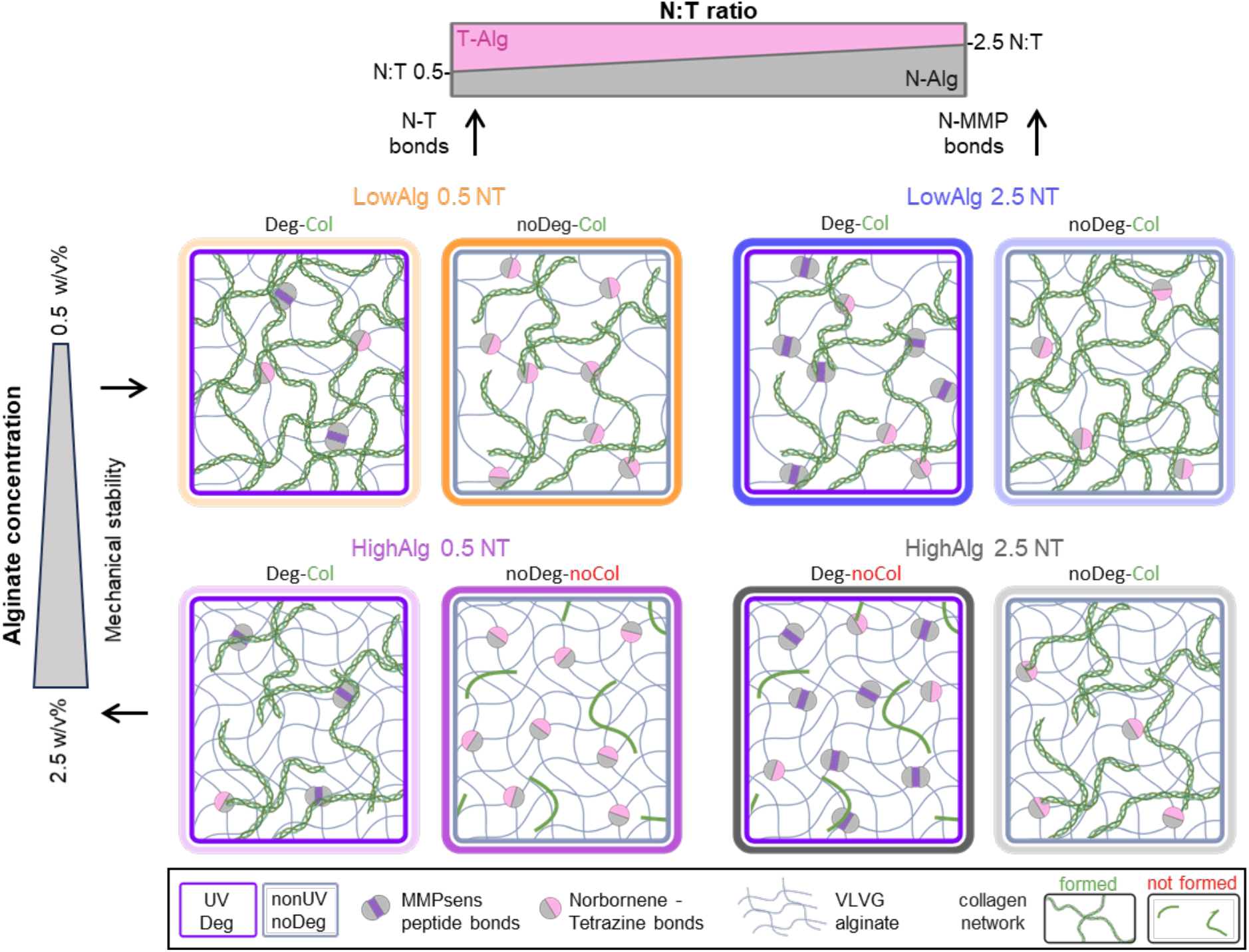
Schematic representation of the Alg-Col IPN material variations. Effect of the alginate N:T crosslinking ratio (0.5 versus 2.5) and alginate concentration (0.5 w/v % versus 2.5 w/v %) on the collagen fiber network formation. Each material can have two variations: N-T bonds (no UV, noDeg, gray rectangle) and MMPsens peptide bonds (UV exposed, Deg, purple rectangle). The color code represents each pair of materials: LowAlg0.5NT (orange), LowAlg 2.5NT (blue), HighAlg 0.5NT (pink) and HighAlg 2.5NT (gray).

For both systems, the precursors for the hydrogel were dissolved in HEPES buffer and distributed into 3 tubes. The first tube contained N-Alg; MMP-sensitive (MMPsens) peptide (GCRD-VPMS ↓ MRGG-DRCG, 95 % purity; WatsonBio) at a final concentration of 5 mg/mL of hydrogel; and thiolated RGD-peptide (CGGGGRGDSP; Peptide2.0) at a final concentration of 1.44 mg/mL of hydrogel (DS 5, 1.2 × 10^-3^ mM). The second tube contained tetrazine-modified alginate (T-Alg) and the photoinitiator (LAP, #900889, Sigma) at a final concentration of 1.5 mg/mL of hydrogel. The third tube contained the collagen precursor solution (10 mg/mL, type I bovine collagen solution, FibriCol 5133, Advanced Biomatrix), used at a final concentration of 5 mg/mL of hydrogel and neutralized prior to the mixing with 10 % HEPES buffer and 10 % NaHCO_3_ (37 mg/mL, pH 9, Sigma, #S6014,). The 3 tubes were mixed and briefly centrifuged to avoid bubble formation. The total final concentration of collagen is constant for all mixtures (5 mg/mL) and the alginate concentration was varied from 0.5 % w/v for low concentration (LowAlg) to 2.5 % w/v for high concentration (HighAlg), at an N:T ratio of 0.5 or 2.5 for each type.

To induce collagen fibrillogenesis, the gels were placed at 37 °C in the dark for 50 min immediately after mixing, and light exposure in the case of the degradable hydrogel, to allow for physical collagen crosslinking simultaneously with the covalent crosslinking. Both fully crosslinked hydrogels were finally exposed to UV light (365 nm) at 5.7 mW/cm^2^ (Omnicure S2000) for 10 s to ensure that the RGD peptide binds to the uncoupled norbornenes via thiol-ene reaction. This step is needed in the case of the non-degradable hydrogels as no thiol-ene click reaction took place and is kept constant for the degradable hydrogels to be compatible with patterning experiments. Hydrogels were punched out and incubated in PBS for mechanical characterization experiments or in cell culture media at 37 °C and 5 % CO_2_ for cell experiments.

### 2.3 Mechanical characterization of degradable or non-degradable bulk IPNs

The materials without a photo mask (the complete gel exposed to UV or noUV) were used as controls for mechanical testing of bulk properties via rheology and stress relaxation measurements under unconfined compression. The bulk materials were produced as previously described in Section 2.2, under unsterile conditions, pipetted and cast onto a bottom glass plate (borosilicate glass, Filafarm, #1000185), with silicon molds (Sahltec-DE, #DIN53505) of 8 mm in diameter and 2 mm thickness, and immediately covered with a glass slide previously treated with SigmaCoat (≥ 99.5 %; Sigma-Aldrich, #SL2) to prevent adhesion to the gel. The cylindrical hydrogels were extracted from the silicon molds and incubated in PBS at 37 °C.

#### 2.3.1. Rheology

Storage and loss modulus of bulk hydrogels were determined with a Discovery-Hybrid- Rheometer (TA Instruments, USA) via frequency sweeps with a parallel plate geometry of 8 mm (PP08, TA Instruments). The frequency sweeps were performed from 0.01 to 10 Hz and at 0.1 % shear strain at RT (n = 3). A humidity chamber was used to avoid hydrogel dehydration.

To obtain the elastic modulus, first the shear modulus (G) was derived from the storage (G’) and loss (G”) modulus using Rubber’s elasticity theory 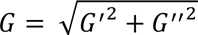 [53]. The elastic modulus (E) was calculated using the values of *G* and the approximation of Poisson’s ratio (P) equal to 0.5, *E* = 2*G* (1 + P) [54].

Additionally, a theoretical approximation of the alginate mesh size is included based on the storage modulus (G’) derived from a previous report [55]. The approximate mesh size (*ξ*) as 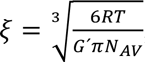, where *R* being the ideal gas constant (8.314 m^3^Pa/Kmol) and *T* being the room temperature (293 K), *G’* the storage modulus and *N_AV_* Avogadrós constant.

#### 2.3.2. Stress relaxation from uniaxial compression testing

The stress relaxation properties of Alg-Col IPN hydrogels were also measured from uniaxial compression testing (HR20, TA Instruments, Germany). The gel disks were compressed to 15 % strain with a deformation rate of 1 mm/min. Subsequently, the strain was held constant, while the load was recorded as a function of time during 1000 s. A humidity chamber was used to avoid hydrogel dehydration. The stress relaxation half time (τ_1/2_) was obtained directly from the graph of normalized force (norm. to force at t = 0) vs. time.

### 2.4. Degradation properties

The degradation of the hydrogels was characterized by monitoring changes in volume, wet weight and dry weight of bulk materials in a collagenase solution (C0130, Sigma). The bulk materials were cast as previously described in Section 2.3, and equilibrated overnight in PBS at 37°C. The initial wet weight of all equilibrated gels was recorded, then the gels were incubated in a collagenase solution (0.01 U/mL) in PBS at 37 °C. Measurements of wet weight after removing the excess liquid with Kimwipe were taken on day 0, 1, 3, 5 and 7.

The hydrogel volume was measured using a caliper. The height was measured once and the diameter was measured from two orthogonal positions and then averaged. The volume was calculated considering a cylindrical shape. Measurements were taken on day 0, 1, 3, 5 and 7.

To measure the dry weight, the excess liquid was removed with a Kimwipe and the gels were lyophilized using the Labconco Freezdryer (FreeZone 4.5, USA). Finally, the dry gel weight was recorded on day 0, 1, 3, 5 and 7 and normalized to the weight on d0.

### 2.5. Patterning of Alg-Col IPN by photolithography

The formation of patterned Alg-Col IPN followed the same procedure as described in Section 2.3, with the addition of a photomask placed on top of the cover glass during the UV mediated thiol-ene coupling of the MMPsens peptide with the N-Alg. The photomask had a pattern of straight lines with 300 μm thickness (UV light blocking sections, non-degradable matrix) placed 500 μm apart (UV light permitting sections, degradable matrix). After the 1.5 min UV exposure through the photomask, an incubation of 50 min at 37 °C followed, for the temperature-dependent collagen fibrillogenesis and N-T bond formation. Finally, the fully crosslinked gels were exposed to 10 s UV light at 5.7 mW/cm^2^ without a photomask for homogeneous binding of the RGD-peptide.

### 2.6. Mechanical characterization via nanoindentation

Nanoindentation measurements were performed on a high-throughput mechanical screening nanoindenter (Pavone, Optics11 Life, Netherlands). The measurements were used to determine the surface elastic modulus and stress relaxation of the distinct regions of patterned materials. For the elastic modulus, each measurement included a minimum of 144 indentations in a 12 x 12 matrix. For the measurements of stress relaxation, the holding time was increased to 300 s after the surface was detected. Patterned hydrogels were prepared as described in Section 2.5 and casted into 8mm in diameter in silicon molds to be fixed in a 6- well plate with a cyanoacrylate-based glue. All measurements were performed using a cantilever-based probe with a spherical tip (49.5 µm radius, 0.38 N/m stiffness, Optics Life) in displacement mode. The Hertzian model was used to determine the effective Young’s modulus *E*_eff_ (kPa) by fitting the load-indentation curves to a maximum indentation depth of 16 % of the tip radius. All measurements were conducted immersed in PBS buffer at RT.

The data was processed automatically using the software provided with the instrument (Pavone Lite Rev 1.2.0).

### 2.7. Confocal reflectance imaging of fibrillar collagen

The collagen fiber network within the bulk and patterned Alg-Col IPNs was imaged in confocal reflectance mode after 24h equilibration in PBS. For this we used a confocal microscope (SP8 Falcon, Leica, Germany), with a 63x 1.2 NA water immersion objective (Leica, Germany) and with 488 nm argon laser illumination. Z-stacks consisting of 30 images with 0.75 μm spacing were obtained.

### 2.8. Scanning electron microscopy (SEM)

Hydrogels were prepared for SEM imaging through a water-ethanol exchange process. The samples were immersed sequentially in ethanol solutions of increasing concentrations, starting with 10 % ethanol and increased in 10 % increments until absolute ethanol was achieved. For each concentration, the hydrogels were incubated under rocking for 5 minutes at RT. Samples were then stored in absolute ethanol at 4 °C until further processing. Next, the samples were dried in a critical point drier (CPD300, Leica EM, Germany) using a 120 min, 2°C/min temperature gradient between 33 to 44°C, 20 cycle protocol. Finally, the samples were gold sputtered (AGB7340, Agar Scientific, UK) and imaged using the JCM- 600 SEM (JEOL GmbH, Germany). SEM images were taken with an accelerating voltage of 15 kV, under high vacuum (320 Bar), a working distance of 10 mm and a spot size of 4.0.

### 2.9. Cell culture: ECs and hMSCs

Human umbilical vein endothelial cells (HUVECs) were obtained from Lonza (#C2519A). The cells were cultured at 37 °C in EGM-2 media (# CC 3162, Lonza) and used in passage 4.

Additional to the HUVECs, bone marrow derived hMSCs were used as supporting cells. hMSCs from a single donor were obtained from RoosterBio (MSC-007) and cultured in RoosterNourish expansion media (K82003, RoosterBio). The cells were cultured at 37 °C and used in passage 5.

The cells were harvested using TryplE (12604013, Gibco), pelleted, and suspended in the corresponding medium at a concentration of 1 × 10^7^ cells/mL. Prior to the cell seeding, the two cell types were mixed in a 1:1 ratio and cultured in the microfluidics system using a 1:1 HUVECs: hMSCs media combination.

#### 2.9.1. Microfluidic system: incorporating patterned Alg-Col IPN and cell culture

Three-lane microfluidic titer plates (OrganoPlates 4003-400B, MIMETAS, Netherlands) were used for all microfluidic cell culture experiments (see sketch in Suppl. Figure S2). The handling of the microfluidic plates was performed according to the manufactureŕs instructions and under sterile conditions. Before hydrogel loading, every observation window (red square Suppl. Figure S2) was filled with 50 µL PBS to provide optical clarity and prevent gel dehydration. A volume of 2.5 µL of the stock solution of the different hydrogels described in Section 2.2 was pipetted into the gel inlet (see Suppl. Figure S2) and crosslinked by inverting the plate, placing a photomask and irradiating UV light as previously described in Section 2.5. To polymerize the collagen, the device was kept in an incubator for at least 30 min at 37°C, 5 % CO_2_ under sterile conditions. The polymerization time was reduced to avoid the dehydration of the hydrogel inside the microfluidics channel.

2μL of the EC-hMSC cell suspension described in the previous section was dispensed into the upper perfusion inlet and the plate was incubated in a vertical position at 37 °C, 5 % CO_2_ for 2 h. After the cells attached to the side of the perfusion channel in contact with the Alg- Col IPN, 50 µL of 1:1 hMSCs:HUVECs medium was added to the perfusion inlet and outlet wells and the plate was placed on an interval rocker platform for continuous perfusion (OrganoFlow S, MIMETAS, Netherlands) at 37 °C, 5 % CO_2_. The rocker was set at a 7- degree inclination and 8 min cycle time. Medium was refreshed three times a week.

#### 2.9.2. Staining of cells inside the microfluidics chip

After 5 days of culture inside the microfluidics chip, EC invasion was analyzed using CD31, DAPI (nucleus) and Phalloidin (Actin) staining. All incubation steps were performed on the MIMETAS rocker at 7-degree inclination and 1 min cycle time and using a volume of 50 µL (inserted in top and bottom inlets) for all staining and washing steps, if not indicated otherwise. The culture media was removed, and the gels were washed twice with PBS (Merck #D8537). Then, the encapsulated cells were fixed in 4 % paraformaldehyde solution (Sigma-Aldrich, #158127) for 45 min at RT, followed by permeabilization with 0.3 % Triton X-100 (Sigma-Aldrich, #11488696) for 15 min, and washed twice with 3 % bovine serum albumin (BSA, Sigma-Aldrich, #A7906) in PBS for 5 min. The primary CD31 antibody (1:1000, Cell Signaling Technology, mAb#77699) was dissolved in PBS containing 0.1 % TritonX-100 and 3 % BSA and 25 μL of this solution were added to each inlet for overnight incubation at 4 °C. The samples were washed 3 times with of 3 % BSA in PBS per inlet/outlet. Then the simultaneous staining of DAPI, phalloidin and secondary antibody was performed using the following concentrations: 0.5 µM DAPI (Thermo Fisher, #S11381), 1 µg/mL of Phalloidin 488 Conjugate (CruzFluor™, #SC363790) and 5 μg/mL of anti-rabbit secondary antibody (AF555, Thermo Fischer, A21428). The staining was performed protected from light, during 4 h at RT under rocking. Final washing was performed twice with 3 % BSA in PBS for 5 min at RT.

A confocal microscope (Leica SP5, Germany) was used to image the stained cells in the microfluidics chips. The lasers used were 405, 488 and 552 nm with a 10x, 0.3 NA, dry objective (Leica, Germany). The imaging was performed as a z-stack of 250 μm at intervals of 5 μm and merging 4 x 2 tiles (width x height) for the complete view of the observation window.

### 2.10. Image analysis

The confocal reflectance images were evaluated using ImageJ software (version 1.54r) [56] and following an image processing as previously described [57]. First the images were segmented and binarized manually to delimit the fibers. The distribution of the total fiber length was plotted using a linear regression. The average fiber length was determined using the reciprocal of the exponential coefficient of a fitted exponential function [57]. Additionally, the collagen formation was evaluated as a percentage of the area covered by the collagen fibers in relation to the total area, determined using the fiber analysis plugin of ImageJ and the area of the field of view, respectively.

Cell infiltration was evaluated considering the number of cell nuclei (DAPI) below the upper border of the mid-channel (refer to Suppl. Figure S2 for the microchip layout). First, each phase of the material was delimited manually based on the bright field image. Then, the nuclei were counted using the count particle plugin of ImageJ. Additionally, the expression of CD31 was quantified relative to DAPI in each section and by subtracting the background from a control sample (hydrogel stained only with the anti-rabbit secondary antibody).

## 3. Results

### 3.1 Alginate crosslinking determines collagen fibrillogenesis and collagen fiber network formation

First, we evaluated the effect of the type and density of alginate crosslinking and degradability (MMPsens and N-T bonds) on collagen fibrillogenesis and collagen fiber network formation, using bulk, non-patterned materials. For this, as described in Figure 1, two alginate concentrations 0.5 w/v % (LowAlg, Figure 2 A-D) and 2.5 w/v % (HighAlg, Figure 2 E-H), using two different N:T ratios (0.5 and 2.5), were explored. In all conditions, MMPsens was added to the precursor solution and the hydrogels were formed with or without UV light. Only in the case of UV exposure, degradable crosslinks with MMPsens are formed with norbornene via thiol-ene click chemistry. Confocal reflectance was used to visualize the collagen network. This technique allows for non-destructive imaging of the microstructure of the IPN, as fibrillar collagen’s refractive index is higher than water and scatters light, in contrast to the highly hydrated structure of most polymers, like alginate [58]. It enables characterizing collagen fibers in length and area [57].

**Figure 2:**
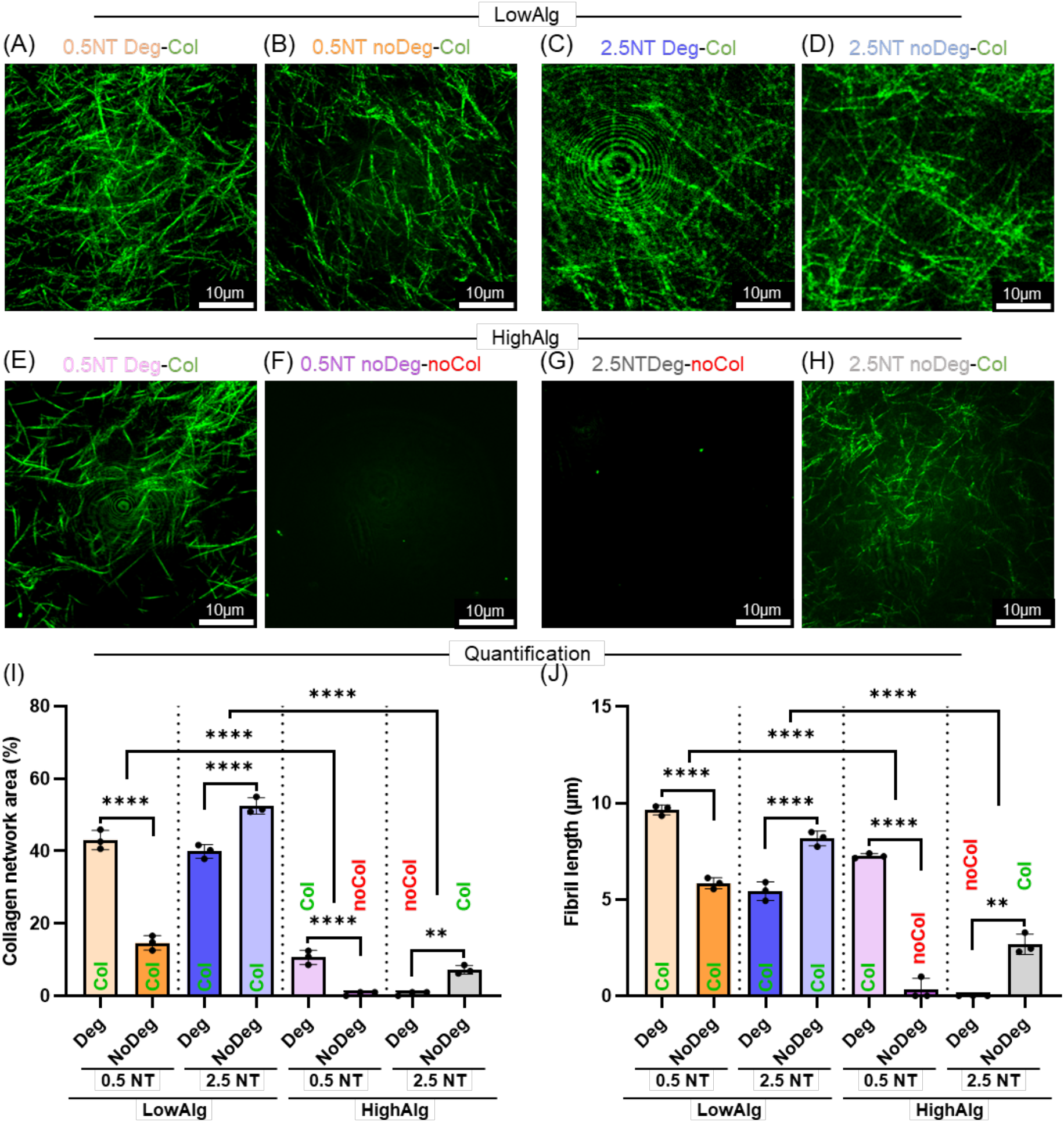
Confocal reflectance of collagen fiber network in bulk IPNs with two different alginate concentrations and two different N:T ratios. Each material variation is composed of the degradable crosslinks (Deg, UV, N-MMPsens crosslink, A, C, E, G) and the non- degradable crosslinks (noDeg, noUV, N-T crosslink, B, D, F, H). The effect of alginate percentage was studied for LowAlg (0.5 w/v %, A-D) and HighAlg (2.5 w/v %, E-H). Different norbornene to tetrazine ratios (N:T) were tested: 0.5NT (A, B, E, F) and 2.5NT (C, D, G, H). The color code in the bar plots indicates the crosslinking level: dark (high) and light (low), and the collagen formation: Col (green) indicates network formed, noCol (red) indicates network not formed. Quantification of the percentage of the collagen area in % (I) and average fibril length in μm (J). Bar plots showing mean and standard deviation of n = 3 different hydrogels. The significant values were evaluated using ordinary two-way ANOVA with Tukey’s correction. Significance indicated with * = p < 0.05, ** = p < 0.01, *** = p < 0.001, **** = p < 0.0001.

Collagen fibrillogenesis occurs at lower crosslinking concentrations. This can be achieved by lower alginate concentration, higher N:T ratios, and higher degradation rate by MMPsens inclusion. For example, the LowAlg 0.5NT Deg-Col, including MMPsens crosslinks, (Figure 2A) has significantly higher collagen area and collagen fiber length compared to its non-Deg equivalent (LowAlg 0.5NT noDeg-noCol, Figure 2B, I, J), based only on NT crosslinks. In contrast, for the 2.5NT ratio, the Deg IPN (LowAlg 2.5NT Deg-noCol, Fig 2C) has a significantly lower collagen area and shorter collagen fiber length when compared to LowAlg 2.5NT noDeg-Col (Figure 2D, I, J). This can be explained by the fact that less NT bonds are formed for 2.5NT compared to 0.5NT but in the presence of light, the excess norbornene will crosslink with MMPsens resulting in more crosslinks for 2.5NT Deg than for 2.5NT noDeg. These results show that even if crosslinks are degradable, the alginate crosslinking density is the main factor determining collagen fibrillogenesis and a collagen network can form in both crosslink types (Deg or noDeg).

In HighAlg (Figure 2 E-H), the differences become even more evident. In the case of HighAlg 0.5 NT Deg-Col (Figure 2E), a collagen network can form, as opposed to HighAlg 0.5NT noDeg-noCol, which does not support any collagen network formation (Figure 2F). The collagen network in HighAlg 0.5 NT Deg-Col differs from its LowAlg equivalent (Fig 2A), with significantly lower collagen fiber area (Figure 2I) and fiber length (Figure 2J) due to the overall increase in crosslinking density. In the HighAlg 0.5NT ratio, stoichiometry reduces the number of norbornene groups available to crosslink with tetrazine or MMPsens. As a result, the HighAlg 0.5NT Deg-Col (Fig 2E) results in a less crosslinked matrix compared to HighAlg 2.5NT Deg-noCol (Figure 2G), enabling collagen fiber network formation (Col), while the latter does not support this, even if both of them are degradable. Interestingly, the high degree of N-T bonds in HighAlg 0.5NT noDeg-noCol material (Figure 2F) hinders collagen fiber network formation, as expected, the HighAlg 2.5NT noDeg-Col (Figure 2H) supports collagen fiber formation, likely due to the lower number of N:T bonds, resulting in a less densely crosslinked and noDeg alginate matrix. These results demonstrate an adverse effect when adding degradable crosslinks in HighAlg 2.5 NT Deg-noCol (Figure 2G), as stoichiometry with excess N increases the number of degradable N-MMPsens bonds, and thus crosslinks in general, resulting in a more crosslinked matrix that does not allow collagen network formation. In contrast, in the HighAlg 2.5NT noDeg-Col (Figure 2H) without UV light, only few N:T bonds can form, resulting in a less densely crosslinked noDeg alginate matrix, enabling collagen fiber network formation. The effect of the N:T ratio on the crosslinking number is supported by a theoretical estimation of the formed bonds of the HighAlg materials, which can be found in Suppl. Table S2.

### 3.2 Collagen fiber network formation modulates elastic and viscoelastic properties

The striking effect of HighAlg in dictating collagen fibrillogenesis established a promising base for further development of anisotropic patterned materials. But first, we investigated the elastic and viscoelastic behavior of bulk, non-patterned HighAlg-Col IPNs using rheology and compression testing followed by stress relaxation measurement (Figure 3). The hydrogels with 0.5NT noDeg-noCol (Figure 3A, dark pink) and 2.5NT Deg-noCol (Figure 3B, dark grey), which have a more densely crosslinked alginate mesh that hinders collagen fiber network formation, show a predominantly elastic behavior with higher storage modulus G’ (Figure 3C), lower loss modulus G” (Figure 3D) and lack of stress relaxing properties (Figure 3E).

**Figure 3:**
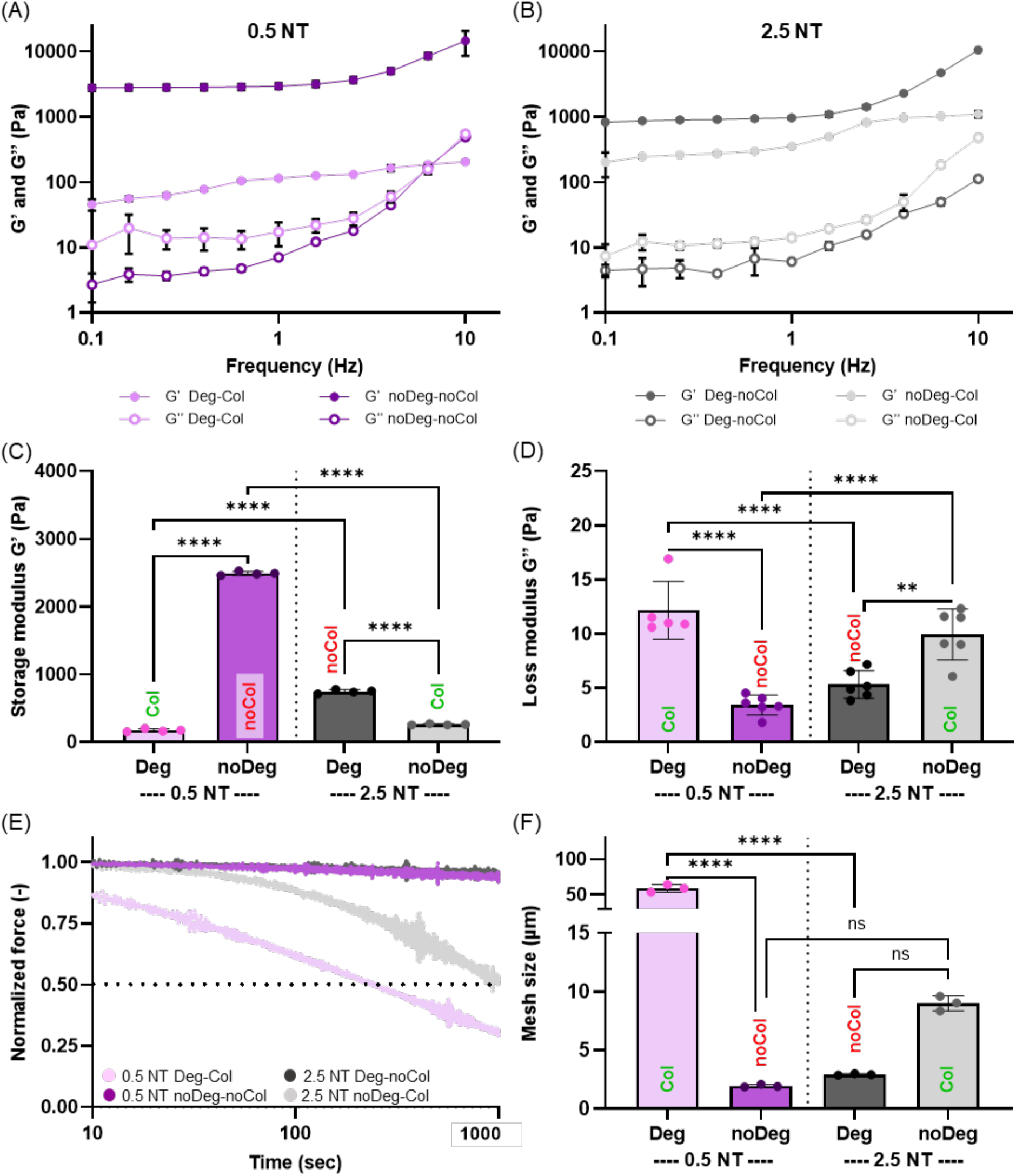
Mechanical characterization of HighAlg IPNs. Storage and loss modulus, G’ and G’’ of bulk materials with 0.5NT (A) and 2.5NT (B) from n = 3. Comparison of G’ (C) and G” (D) in the linear range from n = 3 independent samples. Average stress relaxation of n = 3 hydrogels, depicting the decreasing stress over time for a constant compressive strain of 15 %, normalized to the initial stress (E). Mesh size estimation based on G’ in low frequencies (< 0.1 Hz), from n = 3 independent samples (F). Bar plots showing mean and standard deviation of n = 4 different hydrogels. The significant values were evaluated using ordinary two-way ANOVA with Tukey’s correction. Significance indicated with * = p < 0.05, ** = p < 0.01, *** = p < 0.001, **** = p < 0.0001.

On the other hand, 0.5NT Deg-Col (Figure 3A, light pink) and 2.5NT noDeg-Col (Figure 3B, light grey), which have a looser alginate mesh that enables collagen fiber network formation, show a viscoelastic behavior with predominantly lower storage modulus G’ (Figure 3C), higher loss modulus G” (Figure 3D) and stress relaxing properties (Figure 3E), with a stress relaxation half time of 230 seconds and 950 seconds, respectively. The lower storage modulus G’ for 0.5NT Deg-Col compared to 0.5NT noDeg-noCol suggests that the thiol-ene click chemistry is favored over NT crosslinking when both can happen at the same time.

The mesh size can be theoretically approximated using G’ as previously described [55]. The results show that 0.5NT Deg-Col has the significantly largest estimated mesh size (51.73 ± 3.79 μm), consistent with the lowest estimated crosslinking density (see Suppl. Table S2), and 0.5NT noDeg-noCol has the significantly smallest mesh size (1.48 ± 0.49 μm) consistent with the highest estimated crosslinking density (see Suppl. Table S2).

Based on the previous results, the material properties are summarized in Table 1. In this table, the relationship between the alginate crosslinking density and the formation of the collagen network is detailed.

**Table 1:**
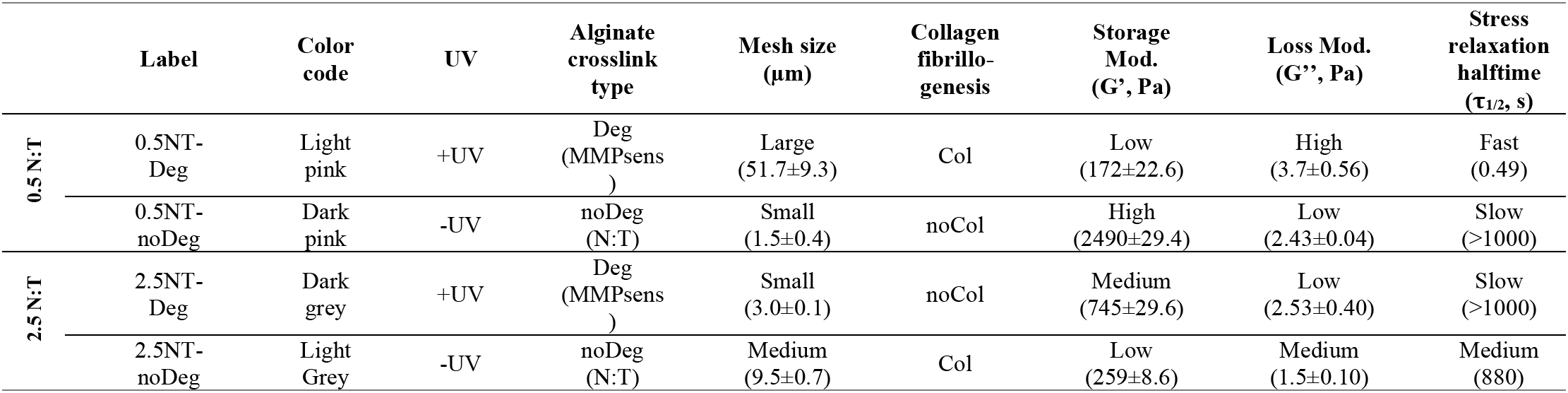
Material combinations of the different HighAlg-Col IPNs and resulting properties in terms of collagen fibrillogenesis, collagen fiber network formation and viscoelasticity.

| | Label | Color code | UV | Alginate crosslink type | Mesh size ($\mu\text{m}$) | Collagen fibrillogenesis | Storage Mod. ($G'$, Pa) | Loss Mod. ($G''$, Pa) | Stress relaxation halftime ($\tau_{1/2}$, s) |
| --- | --- | --- | --- | --- | --- | --- | --- | --- | --- |
| 0.5 N:T | 0.5NT-Deg | Light pink | +UV | Deg (MMPsens) | Large (51.7±9.3) | Col | Low (172±22.6) | High (3.7±0.56) | Fast (0.49) |
| | 0.5NT-noDeg | Dark pink | -UV | noDeg (N:T) | Small (1.5±0.4) | noCol | High (2490±29.4) | Low (2.43±0.04) | Slow (>1000) |
| 2.5 N:T | 2.5NT-Deg | Dark grey | +UV | Deg (MMPsens) | Small (3.0±0.1) | noCol | Medium (745±29.6) | Low (2.53±0.40) | Slow (>1000) |
| | 2.5NT-noDeg | Light Grey | -UV | noDeg (N:T) | Medium (9.5±0.7) | Col | Low (259±8.6) | Medium (1.5±0.10) | Medium (880) |

### 3.3 Alg-Col IPNs mechanical and degradation properties over time

The behavior of these materials also showed different properties over time, as demonstrated by changes in elastic modulus, volume, wet and dry weight over 7 days. The enzymatic degradation of the materials induced by a collagenase solution depends on the alginate crosslink type (Deg or noDeg) and the degree of collagen fiber network formation.

The change in the elastic modulus (Figure 4A) shows the change in stiffness due to the degradation. The elastic modulus of the completely non-degradable material0.5NT noDeg- noCol (dark pink) is stable for at least 7 days, in contrast to the degradable material 0.5NT Deg-Col (light pink), where a steep decline of the stiffness is shown over 7 days. The 2.5NT materials show a gradual decline due to the collagen (2.5NT noDeg-Col, gray) and MMPsens degradable crosslinks (2.5NT Deg-noCol, black) degradation. The noDeg-Col (gray) materials show a fast decline in the elastic modulus due to the degradation of the collagen network, followed by a stabilization by the alginate noDeg bonds. In contrast, the 2.5NT Deg-noCol (black) shows a gradual decline due to the MMPsens degradable bonds. The gels with collagen demonstrate a rapid initial degradation while the MMPsens crosslinks degrades more gradually, likely because collagen is more sensitive to collagenase than MMPsens crosslinks. Furthermore, the 2.5NT noDeg-Col (gray) degradation reaches a plateau as there is a fraction of NT bonds that do not degrade.

**Figure 4:**
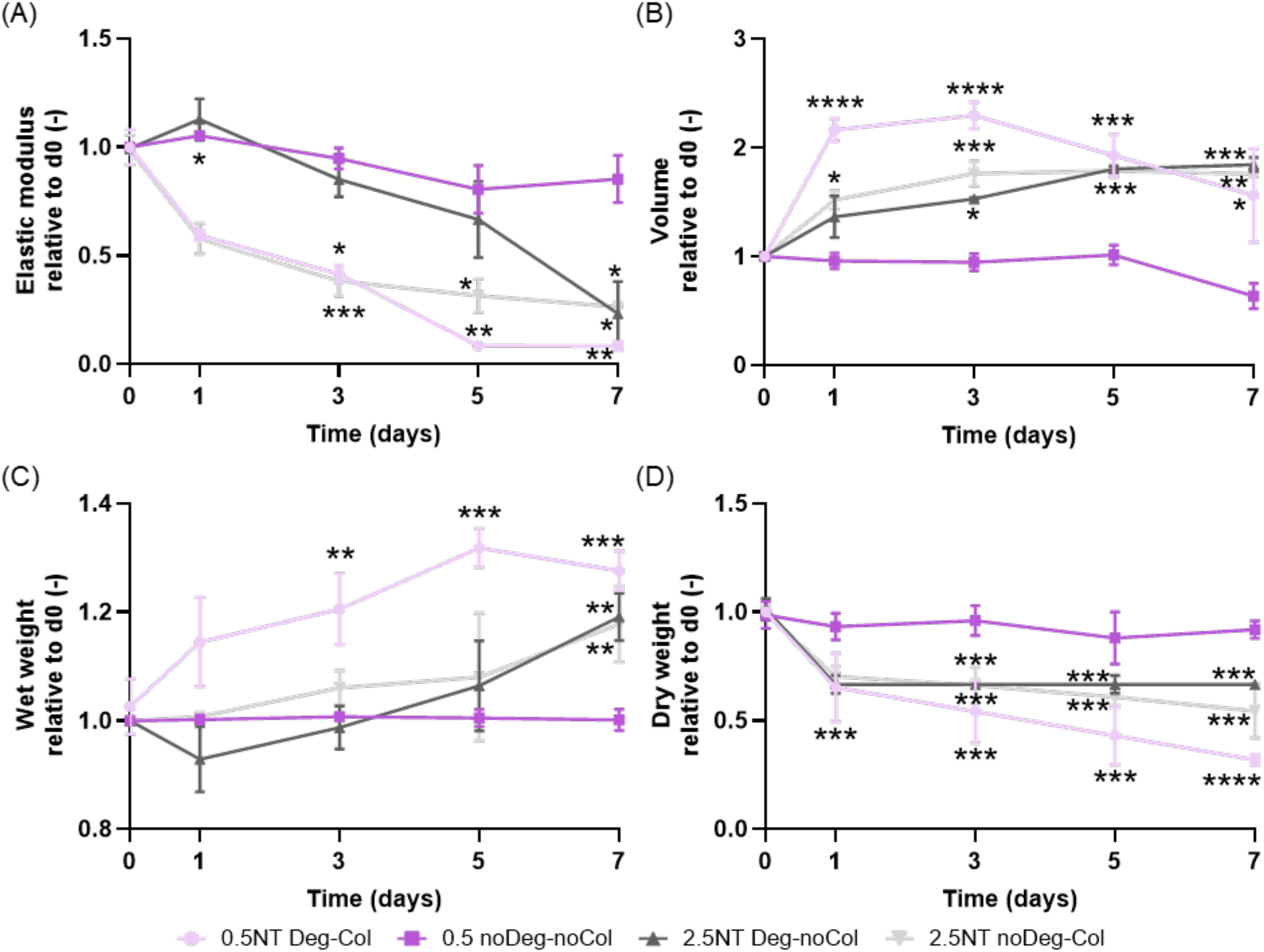
Enzymatic degradation of HighAlg bulk materials incubated with a collagenase solution and tracked over time. Elastic modulus normalized to day 0 (after equilibration in PBS) (A). Volume of the hydrogel disks normalized to d0 (B). Wet weight normalized to d0 (C). Dry weight of hydrogels normalized to the initial dry weight at day 0 (d=5 mm, h=2 mm) (D). Line plots showing mean and standard deviation of time points (1,3,5 and 7 days) of n = 3 gels. The significant differences with respect to d0 were evaluated using ordinary two-way ANOVA with Tukey’s correction. Significance indicated with * = p < 0.05, ** = p < 0.01, *** = p < 0.001, **** = p < 0.0001.

The degradation of the matrix results in water uptake, hydrogel swelling and volume changes (Figure 4B). The lack of degradation of 0.5NT noDeg-noCol (dark pink) is reflected in a stable volume over 7 days, in contrast to 0.5NT Deg-Col (light pink) where a significant increase is shown over the first 4 days, followed by a volume reduction due to degradation. The 2.5NT Deg-noCol (black) material shows a steady change in volume, due to the controlled degradation of the MMPsens peptide crosslinks in the alginate matrix, similar to the degradation of the collagen matrix in a noDeg alginate network of 2.5NT noDeg-Col (grey).

The water uptake that led to the changes in volume can be seen also in the wet-weight changes (Figure 4C). The wet weight of the 0.5NT Deg-Col (light pink) shows a steep increase, in contrast to the constant behaviour of 0.5NT noDeg-noCol. The 2.5NT materials also present a gradual increase in the wet weight over 7 days.

Finally, hydrogel degradation is characterized by measuring the polymer content as dry weight variations over time (Figure 4D). 0.5NT Deg-Col (light pink) shows the greatest decrease in dry weight, in contrast to the 0.5NT noDeg-noCol (dark pink) which shows the most stable behavior over 7 days. In the case of the 2.5NT materials, both show a more gradual decrease in the polymer content due to the degradation of MMPsens degradable bonds (2.5NT Deg-noCol, black) and the collagen network (2.5NT noDeg-Col, gray). The combination of alginate and collagen provides a broad range of materials with differences in their degradation behavior, showing the dynamic nature of the collagen-alginate IPNs.

### 3.4 Patterned IPNs by photolithography

The remarkable differences in properties shown in the bulk HighAlg-Col IPNs can be employed to generate anisotropic 3D patterned IPNs using photolithography. Figure 5 shows the confocal reflectance, SEM and nanoindentation of 3D patterned IPNs by spatially illuminating the previously described bulk materials, 0.5NT (Figure 5A-F) or 2.5NT (Figure 5 G-L).

**Figure 5:**
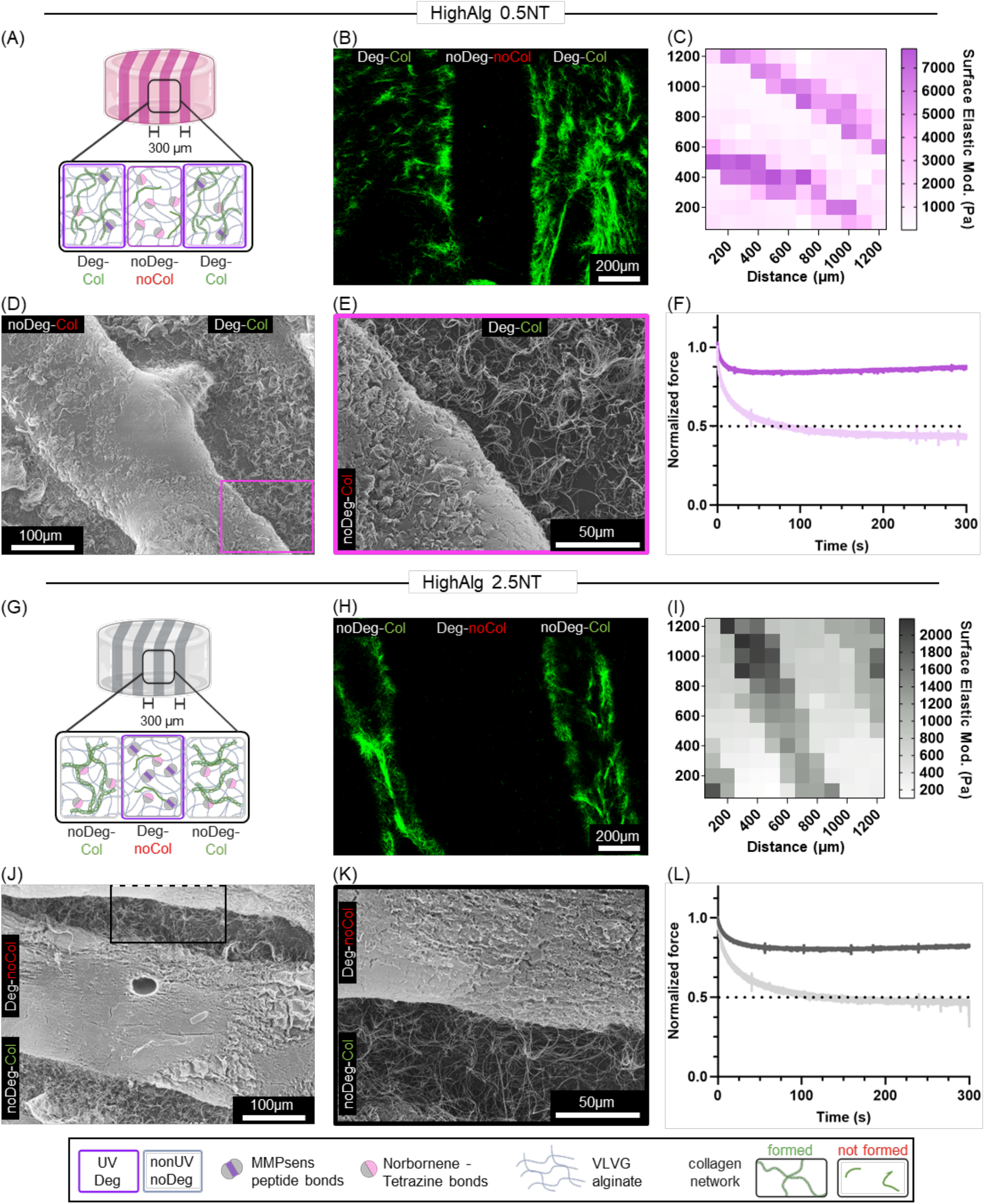
Alg-Col IPNs patterned with photolithography, evaluated using confocal reflectance, SEM and nanoindentation. Sketch of patterned hydrogels and the distribution of the collagen fiber network based on the alginate crosslinks for 0.5NT (A) and 2.5NT (G). Confocal reflectance imaging of patterned materials 0.5NT (B) and 2.5NT (H). SEM images of the surface of patterned materials at 150x (0.5NT (D) and 2.5NT (J)) and 500x (0.5NT (E) and 2.5NT (K). Surface mechanical characterization of collagen-alginate patterned IPNs: Representative nanoindentation of the surface of patterned IPNs yield the surface elastic modulus in a 12 x 12 matrix of 0.5NT (C) and 2.5NT (I) patterned IPNs (n = 1). Averaged single indentations with a holding time of 300 seconds were used to estimate the stress relaxation of the individual phases of 0.5NT (F) and 2.5NT (L) patterned IPNs (n = 3).

The confocal reflectance images (Figure 5B, H) show patterned regions with collagen signal consistent with the softer alginate sections: 0.5NT Deg-Col (Figure 2E) and 2.5NT noDeg- Col (Fig 2H). The impact of the collagen fiber network formation affects the microarchitecture, which can be evaluated using SEM (Figure 5D, E, J, K). The densely crosslinked alginate sections, 0.5NT noDeg-noCol and 2.5NT Deg-noCol, show a more homogeneous surface, compared to the alginate sections with lower crosslinking density, 0.5NT Deg-Col and 2.5NT noDeg-Col (Figure 5E, K). In both cases, a clear interphase is visible between the homogeneous alginate only and the collagen fiber-rich IPN sections (Figure 5D, E, J, K). The characteristic microarchitectures of the patterned materials were evidenced also in the bulk, non-patterned materials (Suppl. Figure S3).

The trends observed in the stiffness and viscoelasticity in the bulk IPNs are consistent with the patterned IPNs. Using nanoindentation, we were able to characterize the elastic moduli and the stress relaxation of the patterned materials (Figure 5C, F, I, L). The IPNs with 0.5NT (Figure 5C) show a pattern in the surface elastic modulus, where the stiffer areas (0.5NT noDeg-noCol, dark pink) have an average elastic modulus of 5.0 ± 1.3 kPa and the softer areas (0.5NT Deg-Col, light pink) 0.5 ± 0.1 kPa. The surface stress relaxation, measured also using nanoindentation in the center of each region, showed differences in the stress relaxation time (Figure 5F), with a viscoelastic region showing a relaxation half time τ_1/2_ of 82 ± 0.2 seconds (0.5NT Deg-col, light pink) and an elastic region (0.5NT noDeg-noCol, dark pink).

For the 2.5NT IPNs (Figure 5G-L), the surface elastic modulus differences are less pronounced and the moduli themselves are lower due to the high N:T ratio. The soft phase averages 0.23 ± 0.07 kPa (2.5NT noDeg-Col, grey) and the stiffer phase has an elastic modulus of 1.5 ± 0.4 kPa (2.5NT Deg-noCol, black). The viscoelastic behavior is present in the regions with collagen network formation (2.5NT noDeg-Col, grey) with t_1/2_ equal to 104 ± 1.2 seconds, whereas the regions without collagen fibers have elastic properties (2.5NT Deg-noCol, black) (Figure 5L). It is important to highlight that, although the trends are consistent with rheology of bulk materials (Figure 3), the actual values obtained using nanoindentation differ, mainly influenced by the differences in surface tension [59], the different nature of measuring technique, and, probably, by the interphases of the anisotropic material.

### 3.5 3D Patterned IPNs in microfluidic chips guide EC cell invasion

EC cell migration and invasion is highly relevant in tissue engineering and regenerative medicine, as well as in disease models. Our 3D patterned IPNs allow tuning three biophysical characteristics that play a role in EC cell invasion: fibrous collagen network architecture, matrix viscoelasticity and degradability. Therefore, we evaluated EC invasion using two combinations of 3D patterned IPNs inside a commercially available microfluidic chip.

Figure 6 shows the invasion of EC cells supported by hMSCs in the different regions of spatially patterned materials as described in Figure 5. After 5 days of culture, the cells proliferated and covered the upper channel (Suppl. Figure S4, Supplementary Movies). Moreover, patterns of EC invasion into specific regions of the 3D patterned Alg-Col IPN were observed, even though RGD was coupled in the entire hydrogel by a rapid post- illumination as described in Section 2.6. In the 0.5NT IPNs (Figure 6A), only the UV-exposed regions (0.5NT Deg-Col) supported EC invasion thanks to the collagen fiber network and softer, viscoelastic and MMP-degradable matrix. In contrast, the 0.5NT noDeg- noCol region does not show any EC invasion, due to the lack of collagen fiber network, and its stiffer, elastic and non-degradable matrix.

**Figure 6:**
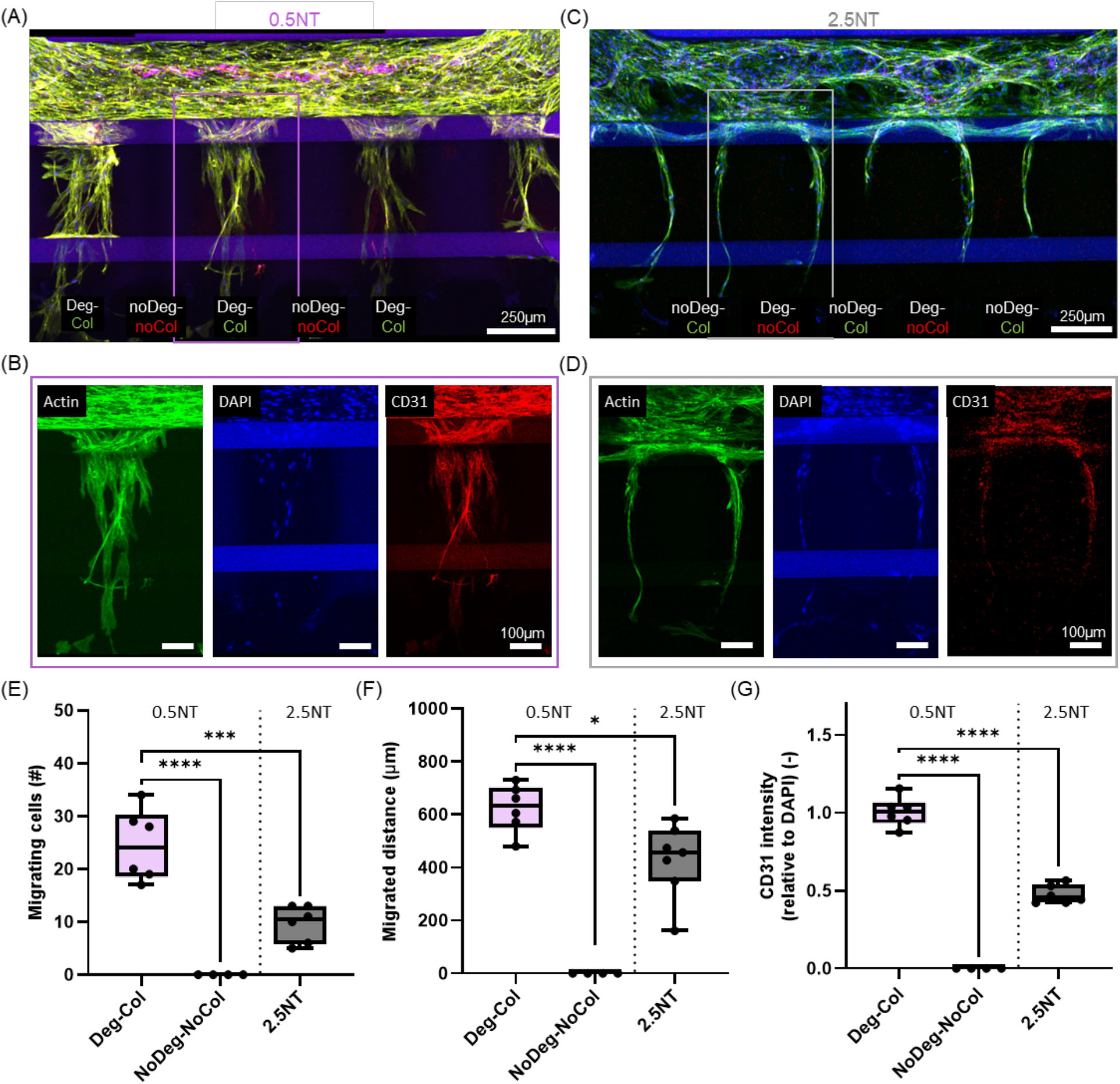
EC invasion supported by hMSCs in 3D patterned Alg-Col IPNs (HighAlg) in a microfluidic chip. Representative maximum projection confocal images of hMSC and EC invasion in patterned materials after 5 days: 0.5NT (A) and 2.5NT (C) stained for DAPI (blue), Phalloidin (green) and CD31 (red). Single channel images are provided for zoom- ins of the hydrogel invaded sections of 0.5NT (B) and 2.5NT (D). Quantification of the invasion distance by ECs (C), cell number (D) and CD31 intensity relative to DAPI (E). Box plots represent the median, max and min of n = 6 phases from n = 3 independent microfluidic chips. Statistical significance with Student t-test for differences between groups is indicated with * = p < 0.05, ** = p < 0.01, *** = p < 0.001, **** = p < 0.0001.

To discern whether EC cell invasion is mediated more by the collagen fiber network or alginate degradability, we further evaluated the 2.5NT materials (Figure 6B). Interestingly, no EC invasion was observed: neither into the 2.5NT Deg-noCol region (no collagen network, elastic, degradable); nor into the 2.5NT noDeg-Col region (rich in collagen, viscoelastic, non-degradable). Instead, the cells take advantage of the interphases between the regions, invading the material. This demonstrates that the MMPsens crosslinking in the 2.5NT Deg-noCol regions is too high and not degrading rapidly enough to enable cell invasion, while the 2.5NT noDeg-Col inhibits cell growth as all crosslinks are not degradable, not providing enough space for cells to infiltrate, even if collagen is present.

Overall, the 0.5NT Deg-Col regions led to the highest EC cell number (Figure 6E) and largest EC cell invasion distance (Figure 6F), compared to their counterpart (0.5NT noDeg-noCol) and to 2.5NT materials. The single channel images reveal higher CD31 expression of ECs in 0.5NT Deg-Col (Figure 6B) materials compared to the cells in the regional interphases of 2.5NT materials (Figure 6D), which is significantly higher when quantified relative to DAPI expression (Figure 6G). Finally, 0.5NT Deg-Col regions when kept for 7 days, resulted in full coverage of the patterned regions and invasion of the lower channel of the microfluidic device (Figure 7).

**Figure 7:**
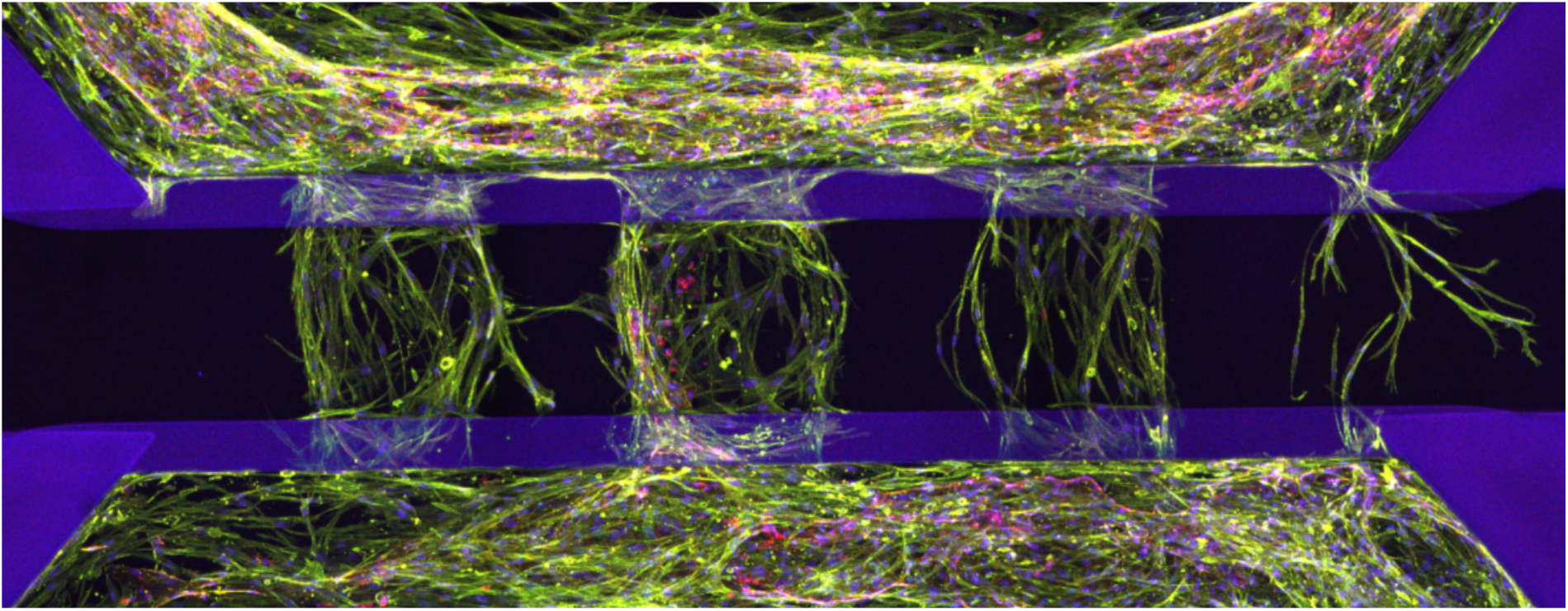
EC invasion supported by hMSCs in 3D patterned Alg-Col IPNs (HighAlg) with 0.5NT after 7 days, stained for DAPI (blue), Phalloidin (green) and CD31 (red).

## 4 Discussion

3D patterned Alg-Col IPNs provide a combination of biophysical properties within a single matrix, including collagen fiber network formation, mechanical (stiffness and viscoelasticity), structural, and degradation properties. While the potential applications of this biomaterial platform are diverse, here we demonstrate that the Alg-Col IPN can direct EC invasion. These results present a novel strategy for developing multicompartment materials by controlling collagen network assembly and enableing the investigation of EC invasion mechanisms.

In Alg-Col IPNs, collagen fibrillogenesis and the resulting collagen fiber network depend strongly on the degree and type of alginate crosslinking (Figure 1). Consistent with previous reports, we found that Alg-Col IPN formation is governed by a competitive crosslinking [35], in which two reactions occur — collagen fibrillogenesis and alginate covalent crosslinking — while mutually interfering with each other’s formation (Figure 2). As a result, the system forms a partially crosslinked structure, often described as a semi-IPN: a network containing only one crosslinked polymer and the chains of the second polymer dispersed in the network [14]. Most Alg-Col IPNs reported to date use only ionically crosslinked alginate [22,23] or sequential crosslinking (ionic followed by covalent), where collagen network formation occurs depending on the dynamic ionic crosslinks of alginate. Our study suggests that collagen assembly can also occur in covalently crosslinked alginate, overcoming limitations associated with the ionic alginate crosslinking strategy (e.g., rapid degradation [29]). This provides a clearer view of collagen–alginate interactions and highlights opportunities for further development of semi-IPN systems.

The degree of covalent bonding in alginate significantly influences the formation of collagen matrix and consequently the resulting mechanical properties, stress relaxation (Figure 3), degradation behavior (Figure 4), and overall microarchitecture (Suppl. Figure 3) of the hydrogel system. Upon formation, the degradable collagen network contributes to viscoelastic properties, attributed to its hierarchical structure and the dynamic nature of hydrogen bond rupture and reformation [60]. In addition, the covalent crosslinking employed here offers two distinct crosslinking mechanisms (MMP-sensitive/degradable and N-T/non-degradable), which provide mechanical stability and preserve structural integrity [27]. In line with prior studies, the resulting IPNs exhibit synergistic behavior arising from emergent physical interactions, interconnected porosity, and a high specific surface area [61,62]. This work demonstrates how controlling the balance between covalent and physical interactions enables the design of IPN hydrogels with tunable degradation kinetics and mechanical performance, making them well-suited for dynamic tissue environments.

Photopatterning of Alg-Col IPNs leads to a multicompartment material that enables spatial segregation of the collagen fiber network, resulting in patterning of mechanical properties and microarchitecture (Figure 5). Previous research on IPNs typically evaluated bulk homogeneous materials and biophysical properties (e.g. stiffness) [39,63]. There are few examples of IPN research to create 3D anisotropic materials: photopatterned HA-collagen multicompartment materials [64], acrylic acid-based IPN (AA-IPN) with controlled deformation [65] and photopatterned bioadhesive collagen-HA [66]. However, in most of these cases, the impact of the material anisotropy is not evident for cell behavior, despite the known advantages of multicompartment materials for guided cell behavior [37,38]. The tunable properties and multicompartment formation of our Alg-Col IPNs provide an ideal platform for guided cell behavior and potential future applications in disease modelling.

The combination of patterned IPNs and organ-on-a-chip technologies offers a platform for assessing cell-material interactions in a physiological relevant scenario. Previous studies showed the relevance of using different matrices or tissue-specific materials, combined with bioprinting, to create 3D structures in multi-organ chips and to predict the interaction between multiple organoids [67,68]. Our photopatterned approach provides an alternative to bioprinting and allows the implementation of cell-guiding structures in microfluidics. On the other hand, this material has great potential to be used as bioink, in combination with light-based technologies capable of creating complex 3D geometries.

Our results underscore the importance of providing collagen such that fiber networks can be formed in degradable materials and EC invasion supported by paracrine signaling from adjacent hMSCs (Figure 6). The adhesion sites provided by the formed collagen network is crucial for ECs and hMSC to interact with degrading matrices, which facilitates cell invasion [69]. In contrast, semi-IPNs formed by a dense alginate matrix and collagen precursors (no collagen fibrillogenesis) hinder cell invasion due to reduced adhesion and an elastic matrix with a small mesh size [70]. However, the availability of adhesion sites is not the sole determinant of EC cell invasion; the matrix’s elastic and viscoelastic properties, as well as its degradability, also play a significant role as they facilitate ECM remodeling by ECs and hMSCs [71]. Consistent with previous research, viscoelastic properties of the ECM can stimulate EC cell invasion [72]. The developed Alg-Col IPN system enables the interplay of these critical characteristics, allowing their combined impacts to be analyzed, providing insights into how biophysical properties — such as the collagen fiber network, elasticity, viscoelasticity, degradation and adhesion — influence EC invasion. This tunable biomaterial tool case can help in tailoring specific cellular niches for future in vitro work where complex cellular microenvironments gain importance.

## 5 Conclusions

The development of 3D IPNs compatible with microfluidics presents a versatile platform for angiogenesis research, tissue engineering and disease models. Here, we propose the use of covalently crosslinked alginate and physically crosslinked collagen to form an IPN. Spatially controlling the alginate crosslinking density and type by photolithography led to patterned collagen fibrillogenesis and consequently, generated anisotropic multicompartment materials with distinct biophysical properties (e.g. microarchitecture, elasticity, viscoelasticity, and degradation properties) to guide cell behavior. Here, such anisotropy in biophysical properties demonstrated the ability to guide EC invasion, supported by hMSCs, in regions with formed fibrillar collagen network, degradable and viscoelastic properties. In contrast, EC invasion was inhibited in phases with non-degradable and elastic or too stiff properties. This work emphasizes the importance of combining several biophysical properties in IPNs to understand cell-material interactions and demonstrates the potential of Alg-Col IPNs for applications in tissue engineering and disease models.

## CRediT authorship contribution statement

C.A. Garrido: Conceptualization, Methodology, Software, Validation, Formal analysis, Investigation, Data Curation, Writing Original Draft, Writing Review & Editing, Visualization. D.S. Garske: Methodology, Software, Validation, Investigation, Data Curation, Writing Original Draft, Writing Review & Editing, Visualization. B. Häßel: Validation, Formal Analysis, Investigation. C. Bastard: Investigation, Writing Review & Editing, Supervision. J. Kamp: Investigation, Writing Review & Editing. L. De Laporte: Supervision, Resources, Writing Review & Editing. K. Schmidt-Bleek & G.N Duda: Supervision, Resources, Writing Review & Editing, Project administration, Funding acquisition. A. Cipitria: Conceptualization, Methodology, Validation, Resources, Writing Original Draft, Writing Review & Editing, Supervision, Project administration, Funding acquisition

## Acknowledgements

The authors acknowledge the support from all group members of Cipitria, Schmidt-Bleek, Duda and De Laporte laboratories. In addition, we thank the Research Workshop at the Charite-Universitätsmedizin Berlin for developing and manufacturing some experimental devices. We thank Andreas Engels for printing the photomasks used in this research.

## Data and code availability

Raw and processed data is available in EDMOND online repository https://doi.org/10.17617/3.QRWOMO.

