## Supplementary Information for "Patterned alginate hydrogel spatially guides collagen fibrillogenesis, viscoelasticity and endothelial cell invasion"

- Supplementary Figure S1: NMR spectra of modified VLVG alginate
- Supplementary Table S1: Degree of substitution of modified alginate
- Supplementary Figure S2: MIMETAS, a commercial microfluidics platform used for endothelial cell invasion assay.
- Supplementary Information S1: Explanation of the considerations for the calculation of the material formation and bonds number.
- Supplementary Table S2: Approximation of bond formation in degradation - patterned hydrogels for HighAlg (2.5 w/v %) and two N:T ratios
- Supplementary Figure S3: SEM images of the surface of single-phase materials.
- Supplementary Movie S1: Videos of 3D reconstruction of the microfluidics chips with HUVEC migration in patterned materials.

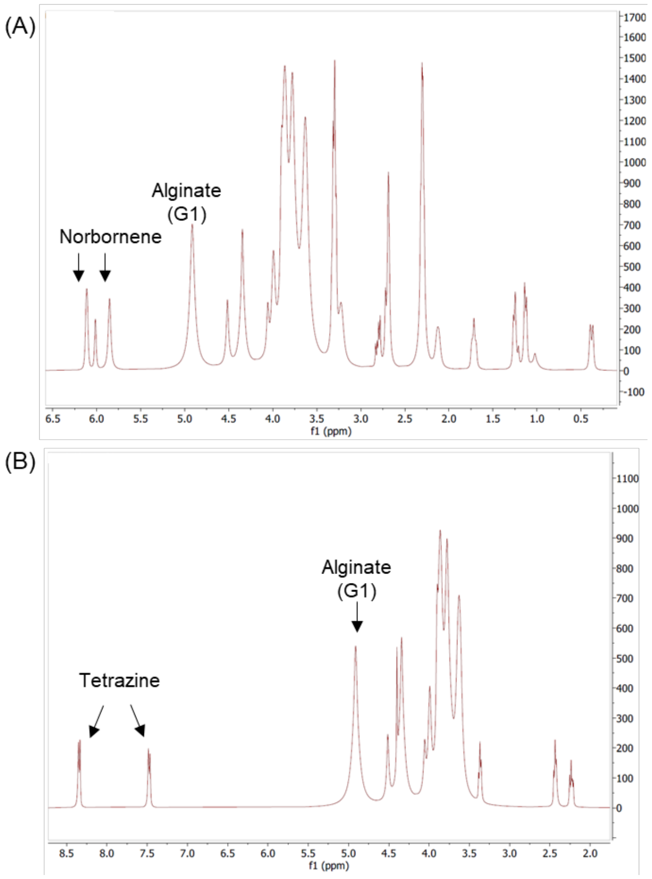


**Supplementary Figure S1: NMR spectra of modified VLVG alginate.** NMR of norbornene modified alginate with the 3 characteristic peaks of norbornene between 6.2-5.8 ppm (A). NMR of tetrazine modified alginate with the 3 characteristic peaks of tetrazine at 10.2, 8.2 and 7.4 ppm (B). In both graphs the H in the first position of the guluronic acid in alginate (G-1), corresponding to the peak between 5.2-4.8 ppm, is used as a reference to determine the $DS_{actual}$.

| | Norbornene | Tetrazine |
|---|---|---|
| $DS_{theo}$ | 500 | 170 |
| $DS_{actual}$ | 32.7 | 33.3 |

**Supplementary Table S1: Degree of substitution of modified alginate.** Norbornene (N-Alg) and tetrazine (T-Alg) modified alginate with a theoretical degree of substitution ($DS_{theo}$) and actual DS ($DS_{actual}$).

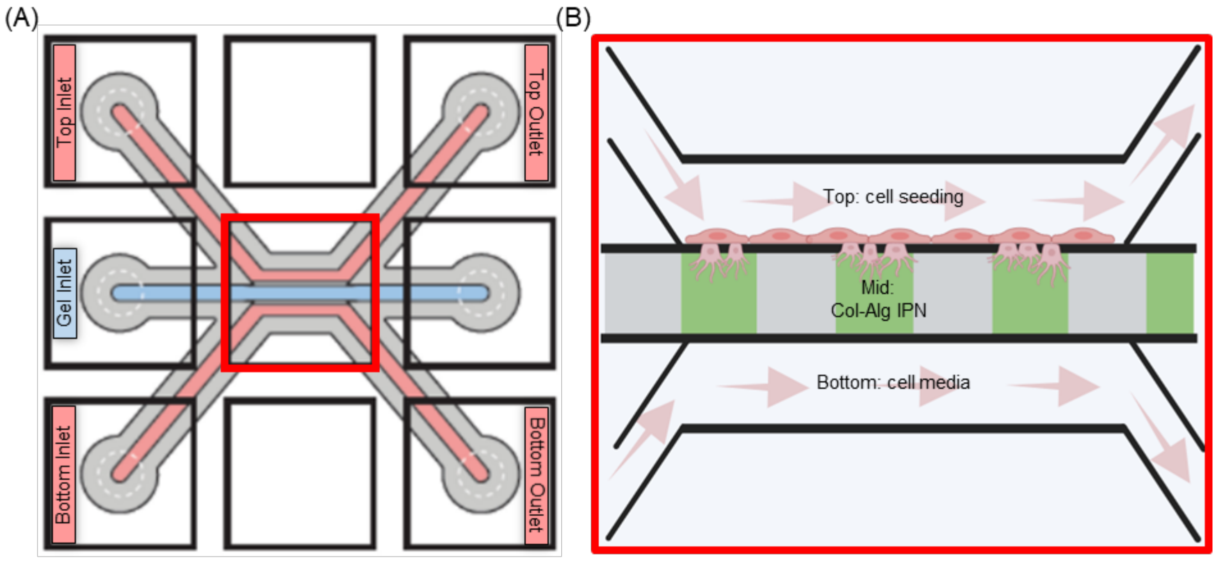


**Supplementary Figure S2: MIMETAS, a commercial microfluidics platform used for endothelial cell invasion assay.** General overview of one of the microfluidics chips and the 6 inlets/outlets (A). Closer view of the observation window and distribution the endothelial cell culture and the Alg-Col IPN (B).

**Supplementary Information S1: Explanation of the considerations for the calculation of the material formation and bonds number.**

The first step is calculating the required mass of N-Alg ($m_N$) and T-Alg ($m_T$), considering the total polymer mass ($m_{Total}$) and $N{:}T\ ratio$ ($RT_{NT}$), determined by the two equations:

$$m_{Total} = m_N + m_T \quad (1)$$

$$R_{NT} = \frac{n_N}{n_T} \quad (2)$$

The number of functional groups ($n$) can be described as a function of the mass ($m$), molecular weight ($MW$) and degree of substation ($DS$) for norbornene (3.1) and for tetrazine (3.2):

$$n_N = \frac{m_N}{MW_N} DS_N \quad (3.1) \qquad n_T = \frac{m_T}{MW_T} DS_T \quad (3.2)$$

If we replace (3.1) and (3.2) in (2) and consider that $MW$ is 75 kDa for both polymers, neglecting the additional weight of functional groups, we have:

$$R_{NT} = \frac{m_N\, DS_N}{m_T\, DS_T} \quad (4) \quad \text{or solved for} \quad m_N = \frac{R_{NT}\, m_T\, DS_T}{DS_N} \quad (4.1)$$

Equation (1) can then be solved for $m_T$ and placed into (4.1):

$$m_N = (m_{Total} - m_N)\frac{R_{NT}\, DS_T}{DS_N} \quad (5)$$

Rearranging and solving equation (5) for $m_N$ results in:

$$m_N\left(\frac{R_{NT}\, DS_T}{DS_N} + 1\right) = m_{Total}\frac{R_{NT}\, DS_T}{DS_N}$$

$$m_N = \frac{m_{Total}\ R_{NT}\ DS_T}{DS_N + R_{NT} DS_T} \quad (6)$$

| | | 0.5NT | | 2.5NT | |
|---|---|---|---|---|---|
| | | Deg | noDeg | Deg | noDeg |
| Reactants | Norbornene (molec/mL) | $2.21 \times 10^{18}$ (1) | $2.21 \times 10^{18}$ | $3.97 \times 10^{18}$ | $3.97 \times 10^{18}$ |
| | Tetrazine (molec/mL) | $4.43 \times 10^{18}$ | $4.43 \times 10^{18}$ | $2.64 \times 10^{18}$ | $2.64 \times 10^{18}$ |
| | MMPsens (molec./mL) | $1{,}78 \times 10^{18}$ (2) | $1{,}78 \times 10^{18}$ | $1.78 \times 10^{18}$ | $1.78 \times 10^{18}$ |
| | RGD (molec/mL) | $7{,}20 \times 10^{17}$ (3) | $7{,}20 \times 10^{17}$ | $7.20 \times 10^{17}$ | $7.20 \times 10^{17}$ |
| Crosslinks | Deg bonds formed | $1{,}11 \times 10^{18}$ (4) | Not formed (5) | $1.78 \times 10^{18}$ | Not formed |
| | N:T bonds formed | Unknown (6) | $2.21 \times 10^{18}$ (5) | Unknown | $2.64 \times 10^{18}$ |
| | RGD coupled | Unknown (6) | Unknown | $4.15 \times 10^{17}$ | $7.20 \times 10^{17}$ |

**Supplementary Table S2: Approximation of bond formation in degradation - patterned hydrogels for HighAlg (2.5 w/v %) and two N:T ratios**

(1) Sample calculation of the amount of norbornene functional groups considering the DS values of Supplementary Table S1, 2.5 % gel and 0.5 N:T ratio in 1mL, we require 8.4 mg of N-Alg and 16.5 mg of T-Alg. Obtained using the equation (6) derived in Supplementary Information S1.

$$8.4mg\ of\ N\ Alg\ \frac{1g\ N\ Alg}{1000\ mg\ N\ Alg} \times \frac{1M\ N\ Alg}{75000\ g\ N Alg} \times \frac{6.022 \times 10^{23}\ molec.N\ Alg}{1M\ N\ Alg} \times \frac{32.7\ N\ groups}{1\ molec.N\ Alg} = 2{,}21 \times 10^{18}\ N\ groups$$

(2) MMPsens peptide is used at a final concentration of 5 mg/mL gel solution, with a MW of 1696 g/mol.

(3) RGD is used at a final concentration of 1.03 mg/ml gel solution, with a MW of 861.89g/mol.

(4) In Deg regions UV-initiated thiol-ene bond formation occurs first and uses up all available MMPsens molecules at an MMP:norbornene stoichiometry of 1:2. This calculation considers the maximum of degradable crosslinks possible.

(5) In noDeg, spontaneous Diels-Alder bond formation uses up all available tetrazine molecules (limiting reagent), therefore there are no Deg bonds formed.

(6) Our approximation simplifies the degradable bond calculation by disregarding the complex simultaneous reactions: the spontaneous N-T crosslinks formation and the coupling of RGD during the UV exposure.

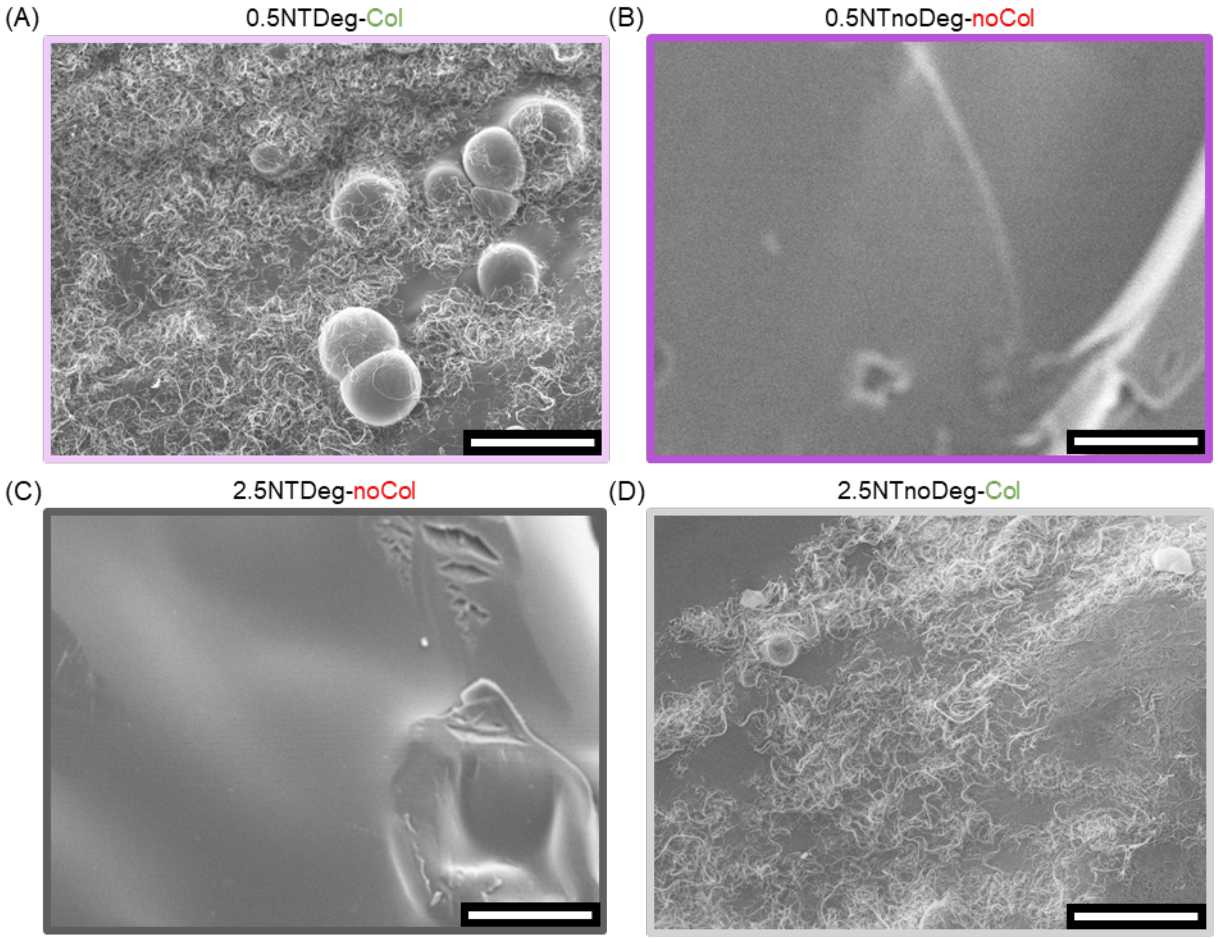


**Supplementary Figure S3: SEM images of the surface of bulk HighAlg materials at 240x**. 0.5NT Deg-Col (A) and 0.5NT noDeg-noCol (B) and 2.5NT Deg-noCol (C) and 2.5NT noDeg-Col (D).

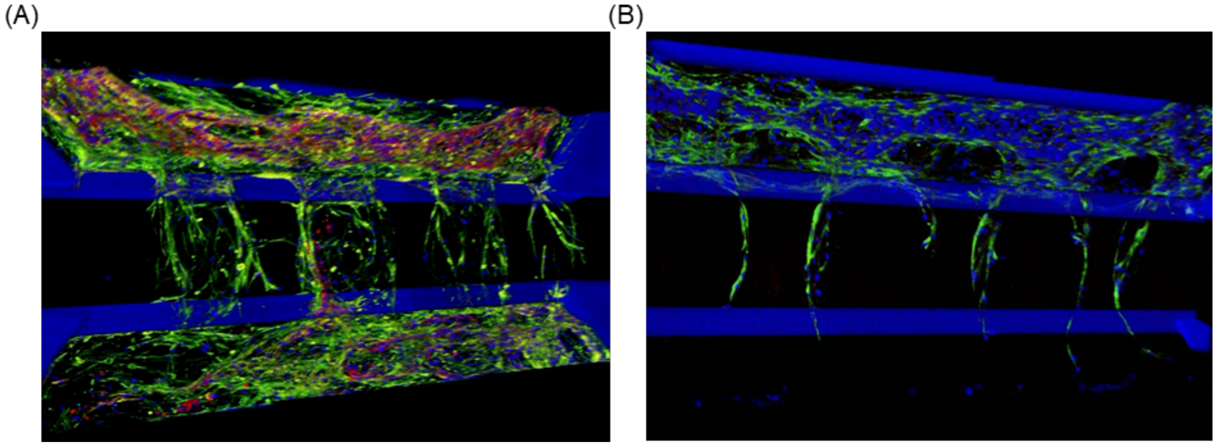

**Supplementary Figure S4: Videos of 3D reconstruction of the microfluidics chips with HUVEC-hMSC invasion in 3D patterned Alg-Col IPNs.** Representative confocal images of hMSC and EC invasion in patterned materials: 0.5NT after 7 days (A) and 2.5NT after 5 days (B) stained for DAPI (blue), Phalloidin (green) and CD31 (red). The 3D reconstruction was performed using LAS X software (v. 3.10, Leica).
